# Dynamic Allostery Highlights the Evolutionary Differences between the CoV-1 and CoV-2 Main Proteases

**DOI:** 10.1101/2021.10.02.462863

**Authors:** P. Campitelli, J. Lu, S. B. Ozkan

## Abstract

The SARS-CoV-2 coronavirus has become one of the most immediate and widely-studied systems since its identification and subsequent global outbreak from 2019-2020. In an effort to understand the biophysical changes as a result of mutations, the mechanics of multiple different proteins within the SARS-CoV-2 virus have been studied and compared with SARS-CoV-1. Focusing on the main protease (mPro), we first explored the long range dynamic-relationship, particularly in cross-chain dynamics, using the Dynamic Coupling Index (DCI) to investigate the dynamic coupling between the catalytic site residues and the rest of the protein, both inter and intra chain for the CoV-1 and CoV-2 mPro. We found that there is significant cross-chain coupling between these active sites and distal residues in the CoV-2 mPro but it was missing in CoV-1. The enhanced long distance interactions, particularly between the two chains, suggest subsequently enhanced cooperativity for CoV-2. A further comparative analysis of the dynamic flexibility using the Dynamic Flexibility Index (DFI) between the CoV-1 and CoV-2 mPros shows that the inhibitor binding near active sites induces change in flexibility to a distal region of the protein, opposite in behavior between the two systems; this region becomes *more* flexible upon inhibitor binding in CoV-1 while it becomes *less* flexible in the CoV-2 mPro. Upon inspection, we show that, on average, the dynamic flexibility of the sites substituted from CoV-1 to CoV-2 changes significantly less than the average calculated across all residues within the structure, indicating that the differences in behaviors between the two systems is likely the result of allosteric influence, where the new substitutions in COV-2 induce flexibility and dynamical changes elsewhere in the structure.

**SIGNIFICANCE:** Here we have conducted a comparative analysis between the SARS-CoV-1 and SARS-CoV-2 mPro systems to shed mechanistic insight on the biophysical changes associated with the mutations between these two enzymes. Our work shows that the CoV-2 mPro system exhibits enhanced cross-chain communication between catalytic site residues and the rest of the structure. Further, both dynamic coupling and dynamic flexibility analyses indicates that, largely, the dynamic changes as evaluated by DCI and DFI occur at sites *other than* the mutation sites themselves, indicating that the functional differences between these two proteins are a result of dynamic allostery

## INTRODUCTION

From its initial onset in late 2019, SARS-CoV-2 or the coronavirus-2, associated with acute respiratory disease, arose in Wuhan, China. Spreading rapidly through over 200 countries resulting in millions of deaths worldwide, the SARS-CoV-2 quickly became the cause of an unprecedented global pandemic in the modern era. In a massive effort to combat the viral spread, effects and subsequent toll on human health and life, large swathes of the scientific community including disciplines ranging from genetics, evolutionary biology, biological physics, data science, immunology and others have taken part in a focused effort to stymie the contagion outbreak resulting in an unparalleled development and production of vaccinations within one year of the virus’ discovery. In December of 2020, the U.S. Food and Drug Administration issued the emergency use authorization (EUA) for the Pfizer BioNTech(1) and Moderna(2) vaccines.

Positing ∼95% prevention of SARS-CoV-2 infection(3,4), these vaccinations presented themselves as enormous breakthroughs in the coronavirus-2 relief effort. However, even with the development of effective vaccinations there still remains a need to continue investigation for further potential SARS-CoV-2 inhibitors. In several studies conducted by experts in the public health sector as well as the scientific community, concerns were raised regarding the overall efficacy of the vaccination process with challenges ranging from logistics and distribution to developmental costs as well as public perception and hesitancy in willingness to undergo the vaccination procedure(5–8). Moreover, SARS-CoV-2 mutation rate in combination with the high level of transmission has resulted in the generation of new, uniquely identifiable strains of the virus including one particular variant of concern, the UK B.1.1.7 lineage which exhibits a higher rate of transmission than other strains with a greater level of interaction with host cell receptors including epithelial cell ACE2(9–12).

Given the high rate of infection, profound global impact of the coronavirus-2 disease and the implication that the virus will continue to go through mutations which may result in more transmissible strains that may prove resistant to currently approved immunization procedures, the continued investigation into not only additional vaccination or treatment methods remain critical. At the heart of further drug discovery, to truly combat and inhibit the virus, an understanding of the virus’ biophysical behavior is required. Particularly, it is important to obtain mechanistic insights into critical proteins of the virus which regulate its ability to interact with host cells and successfully self-replicate for two major reasons. First, to find or design novel drugs or possible allosteric inhibitors, the dynamical behavior of viral proteins must be understood. Second, the mechanical behavior of these proteins and, subsequently, how the shape of the mutational landscape regulates protein dynamics is necessary to understand whether observed mutations will confer resistance to developed drugs.

To that end, we focus on the SARS-CoV-2 main protease (mPro), an enzyme critical for the successful reproduction of the virus upon host cell infection. The mPro is responsible for the processing of two major polyproteins into several nonstructural proteins ultimately responsible for the production of structural proteins comprising the envelope, membrane, spike and nucleocapsid structural proteins(13,14). Thus, the mPro has undergone significant investigation as a potential drug target(15–18). In a comparative analysis between the CoV-1 and CoV-2 mPro enzymes, here we employ two metrics, the Dynamic Flexibility Index (DFI) and the Dynamic Coupling Index (DCI) to evaluate the dynamics and site-specific interactions between different regions of the two proteins. The DFI parameter measures each position’s sensitivity to perturbations within a network of interactions and represents a given amino acid position’s ability to explore its local conformational space (flexibility) while DCI measures the displacement response of an individual position to the perturbation of a second position or group of positions, relative to the average response to any perturbation of all possible positions and can capture the dynamic coupling between amino acid pairs.

When the DCI metric was applied to these two systems, we first found that the cross-chain dynamic coupling is enhanced for the CoV-2 mPro catalytic site residues as compared to the CoV-1 mPro system. The DCI profiles indicated that this type of cross-chain communication is likely an important mechanistic regulator and may be a critical functional difference between these two systems. To further understand the mechanistic changes associated with the virus’ functional evolution we utilized DFI to analyze the flexibility differences and found that, surprisingly, most of the large changes in amino acid flexibility in occurred CoV-2 at sites *other than* the sites that are substituted when CoV-1 and CoV-2 sequences are compared. That is, the mutations brought about dynamical changes in the SARS-CoV-2 mPro at locations distal to the sites of mutation suggesting that allosteric regulation may be a key component in capturing the changes in dynamics between CoV-1 and CoV-2.

To study this behavior further, we next wanted to determine how the allosteric response to ligand binding events (particularly inhibitor binding) differed between the two by incorporating the inhibitor interactions at the active site using the inhibitor bound structures. Here, the CoV-1, mPro (PDB ID 3TIU) and CoV-2 mPro (PDB ID 7BUY) structures were complexed with inhibitors bound to their respective catalytic sites, where four alpha carbons among each inhibitor were used as additional nodes in the ENM network. From these models, we analyzed the structural dynamics associated with ligand-bound mPro forms. Our results showed that dynamic flexibility differences were observed for a set of residues located at the dimeric interface of both chains while the DFI profiles of the rest were largely unchanged between CoV-1 and CoV-2 mPros. Notably, when bound to an inhibitor, these residues of the CoV-1 mPro became *more* flexible, whereas the same residues in the inhibitor-bound CoV-2 mPro became *less* flexible suggesting that interdomain interactions critical for mPro activity(19–21) get much less affected in CoV-2 when inhibitor is bound to active site.

## MATERIALS / METHODS

### Molecular Dynamics Modeling

Topology files for all structures were prepared using the AMBER LEaP program with the ff14SB force field (30). Hydrogen atoms were added and each structure was surrounded by a 16.0 Å cubic box of water molecules using the TIP3P (53) water model. Na^+^ and Cl^-^ atoms were added for neutralization. Each system was energy-minimized using the AMBER SANDER package (30) to remove any unfavorable torsional angles or steric clashes and ensure that the system reached a local energetic minimum. First, the protein was kept fixed with harmonic restraints to allow surrounding water molecules and ions to relax, followed by a second minimization step in which the restraints were removed and the protein-solution was further minimized. Both minimization steps employed the method of steepest descent followed by conjugate gradient. The systems were then heated from 0K to 300K over 250 ps, at which point long-range electrostatic interactions were calculated using the particle mesh Ewald method (54). Direct-sum, non-bonded interactions were cut off at distances of 9.0 Å or greater. The systems were then simulated using Molecular Dynamics at constant temperature and pressure with 2fs time steps for 500 ns. Covariance matrix data were calculated over the final 200 ns of each simulation, using 50 ns moving windows that overlap by 25 ns. To ensure the robustness of our analysis, we used two apo PDB structures separately for both CoV-1(2GZ9, 1UK3) and CoV-2(6Y84, 5R7Y) and conducted the simulations above.

### Dynamic Flexibility Index

The Dynamic Flexibility Index utilizes a PRS technique that combines ENM and LRT (26,23).

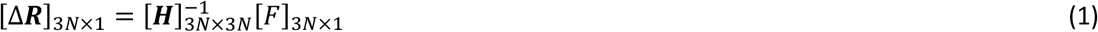

Using LRT, Δ***R*** is calculated as the fluctuation response vector of residue *j* as a result of unit force’s ***F*** perturbation on residue *i*, averaged over multiple unit force directions to simulate an isotropic perturbation.. ***H*** is the Hessian, a 3N × 3N matrix which can be constructed from 3-D atomic coordinate information and is composed of the second derivatives of the harmonic potential with respect to the components of the position’s vectors of length *N*. For this work, the Hessian matrix was extracted directly from molecular dynamics simulations as the inverse of the covariance matrix. This method allows one to implicitly capture specific physiochemical properties and more accurate residue-residue interactions via atomistic force fields and subsequent all-atom simulation data.

Each position in the structure was perturbed sequentially to generate a Perturbation Response Matrix ***A***

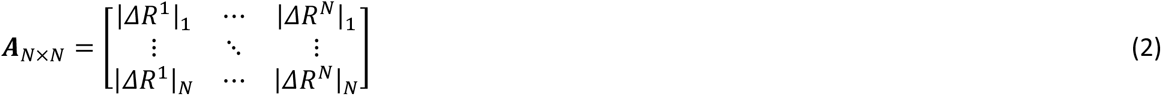

where 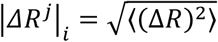 is the magnitude of fluctuation response at position *i* due to perturbations at position *j*. The DFI value of position *i* is then treated as the displacement response of position *i* relative to the net displacement response of the entire protein, which is calculated by sequentially perturbing each position in the structure (3).

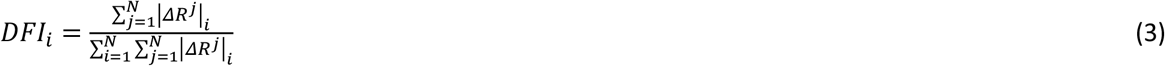

It is also often useful to quantify position flexibility relative to the flexibility ranges unique to individual structures. To that end, DFI can be presented as a percentile rank

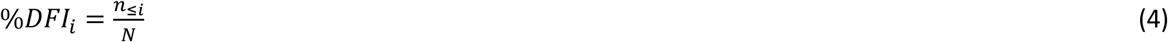

where *n*_*≤i*_ is the number of positions with a DFI value ≤ DFI_i_. The denominator is the total displacement of all residues, used as a normalizing factor. All %DFI calculations present in this work used the DFI value of every residue of the full LacI structure for ranking. The DFI parameter can be considered a measure of a given amino acid position’s ability to explore its local conformational space.

### Dynamic Coupling Index

Similar to DFI, the dynamic coupling index (DCI) also utilizes Perturbation Response Scanning with the Elastic Network Model and Linear Response Theory. The dynamic coupling index (DCI) captures the strength of displacement response of a given position *i* upon perturbation to a single functionally important position (or subset of positions) *j*, relative to the average fluctuation response of position *i* when all of the positions within a structure are perturbed.

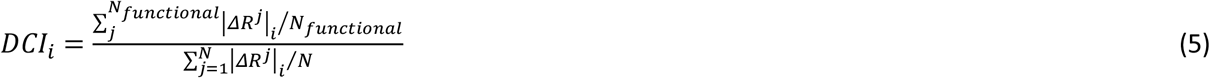

As above, DCI can also be presented as a percentile rank as

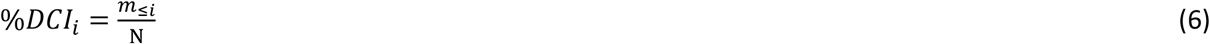

where *m*_*≤i*_ is the number of positions with a DCI value ≤ DCI_i_. As such, this parameter represents a measure of the dynamic coupling between *i* upon a perturbation to *j*.

One of the most important aspects of both DFI and DCI is that the entire network of interactions is explicitly included in subsequent calculations without the need of dimensionality reduction such as Normal Mode Analysis through principal component analysis. If one considers interactions such as allostery as an emergent property of an anisotropic interaction network, it is critical to include the interactions of the entire network to accurately model the effect one residue can have on another.

## RESULTS / DISCUSSION

### Cross-chain Dynamic Coupling is Enhanced for the CoV-2 mPro Catalytic Site Residues

Previous work on the CoV-1 and CoV-2 mPros has indicated the importance of cross-chain communication between chains A and B for function. A previous study conducted by Tom McCleish(22) employed the ENM model using low-frequency modes to study allosteric interactions and dynamics comparing the two proteases. This work showed that several regions of the protein exhibited strong cross-chain dynamic coupling as measured by a residue-residue dynamic cross-correlation map. Further, their work suggested that there are several residues located on the dimeric interface critical to allosteric interactions with the CoV-2 mPro catalytic sites and that there are additional allosteric sites that can significantly change the active site coupling by slight stiffening of local harmonic restraints employed within the ENM model. In the work presented here, we wanted to further investigate the ability for these catalytic sites to allosterically interact with the rest of the structure and, potentially, identify additional putative allosteric residues. Here we implement one of our tools which can determine the strength of dynamic coupling between two residues *i* and *j*, the Dynamic Coupling Index (DCI). Utilizing Linear Response Theory (LRT) and Perturbation Response Scanning (PRS), DCI models mutations or amino acid interactions as fluctuation responses to force perturbations. Specifically, DCI_ij_ measures the displacement response of a given position *j* upon a force perturbation (averaged over many unit force directions to simulate an isotropic Brownian kick) to a given position *i*, normalized by the average response of position *j* when all residues in the protein are perturbed. DCI has been used previously in many different systems to identify important sites of regulation, particularly distal to active site residues named dynamic allosteric residue coupling (DARC) sites which control the dynamics of the active site through dynamic allostery;. particularly mutation of DARC sites could alter function(23–29).

First, we conducted equilibrated molecular dynamics simulations using AMBER ff14SB (30) (see Methods for simulation details) and extracted covariance matrix data taken in moving 50ns windows with a 25ns overlap over the final 200ns of the simulations. We then analyzed the dynamic coupling via averaged values from each covariance matrix between the catalytic residues H41 and C145 of chain A and the rest of the structure by applying force perturbations to H41 and C145 simultaneously, the obtained DCI profiles are then rescaled using percentile rankings (i.e. %DCI). Fig. 1 shows the graphical depiction of this analysis.

**Figure 1.**
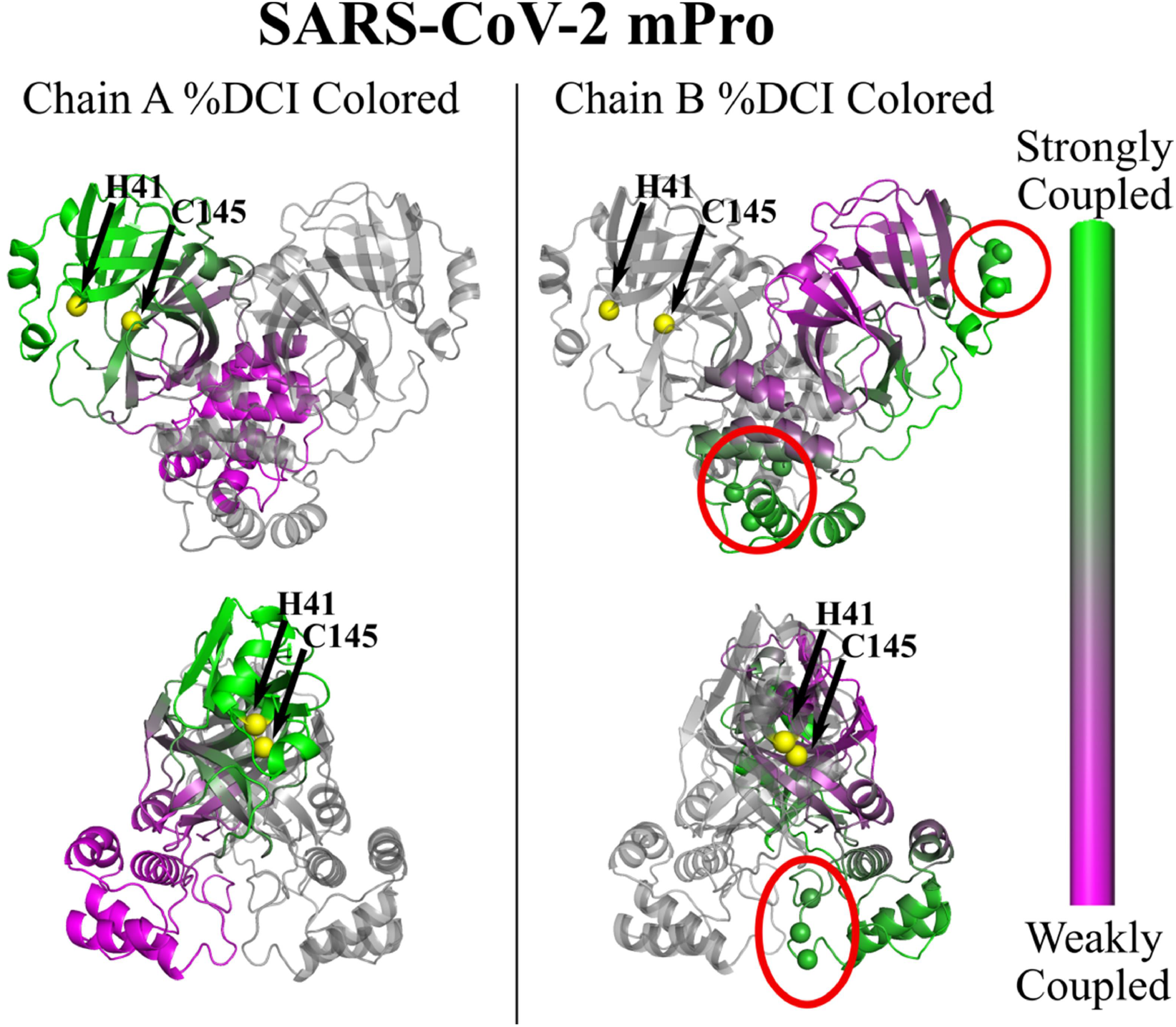
Dynamic coupling analysis (measured by %DCI) between catalytic site residues of H41 and C145 of chain A (yellow spheres) and the other residues of chain A (left) and chain B (right) mapped onto the SARS-CoV-2 mPro (PDB ID 5R7Y(31), inhibitors removed). Green regions of each structure indicate stronger dynamic coupling while purple represents weak dynamic coupling. Within the chain (left), the %DCI distribution is as expected based off proximity; residues close to the catalytic sites are strongly coupled, with those further away exhibiting weak coupling. However, for A-to-B interactions (right), this behavior is nearly the opposite of what one would expect if coupling was a measure of proximity alone. Here, regions of chain B close in space to the chain A catalytic sites are weakly coupled to these sites, whereas some distal portions are very strongly coupled. Chain B sites which exhibited particularly strong coupling to the chain A catalytic residues are circled in red and represented as spheres, comprised of residues E55, I59 and R60 (top right, top circle) and residues N277, R279 and L286 (top and bottom right, bottom circle).

Within a given chain, the %DCI distribution is, as expected, based off proximity; residues close to the catalytic sites are strongly coupled, with those further away exhibiting weak coupling. However, for chain A-to-B interactions, this behavior is nearly the opposite of what one would expect if coupling was a measure of proximity alone. Here, regions of chain B close in space to the chain A catalytic sites are weakly coupled to these sites, whereas some distal portions are very strongly coupled. Chain B sites which exhibited particularly strong coupling to the chain A catalytic residues are circled in red, comprised of residues E55, I59 and R60 and residues N277, R279 and L286.

These coupling profiles further indicate that cross-chain communication is likely an important mechanistic regulator for the proper functioning of the SARS-CoV-2 mPro. In fact, a direct comparison of the same analysis to the CoV-1 mPro shows that this strong inter-chain coupling between the catalytic sites and these residue groups is nearly lost completely (Fig. 2).

**Figure 2.**
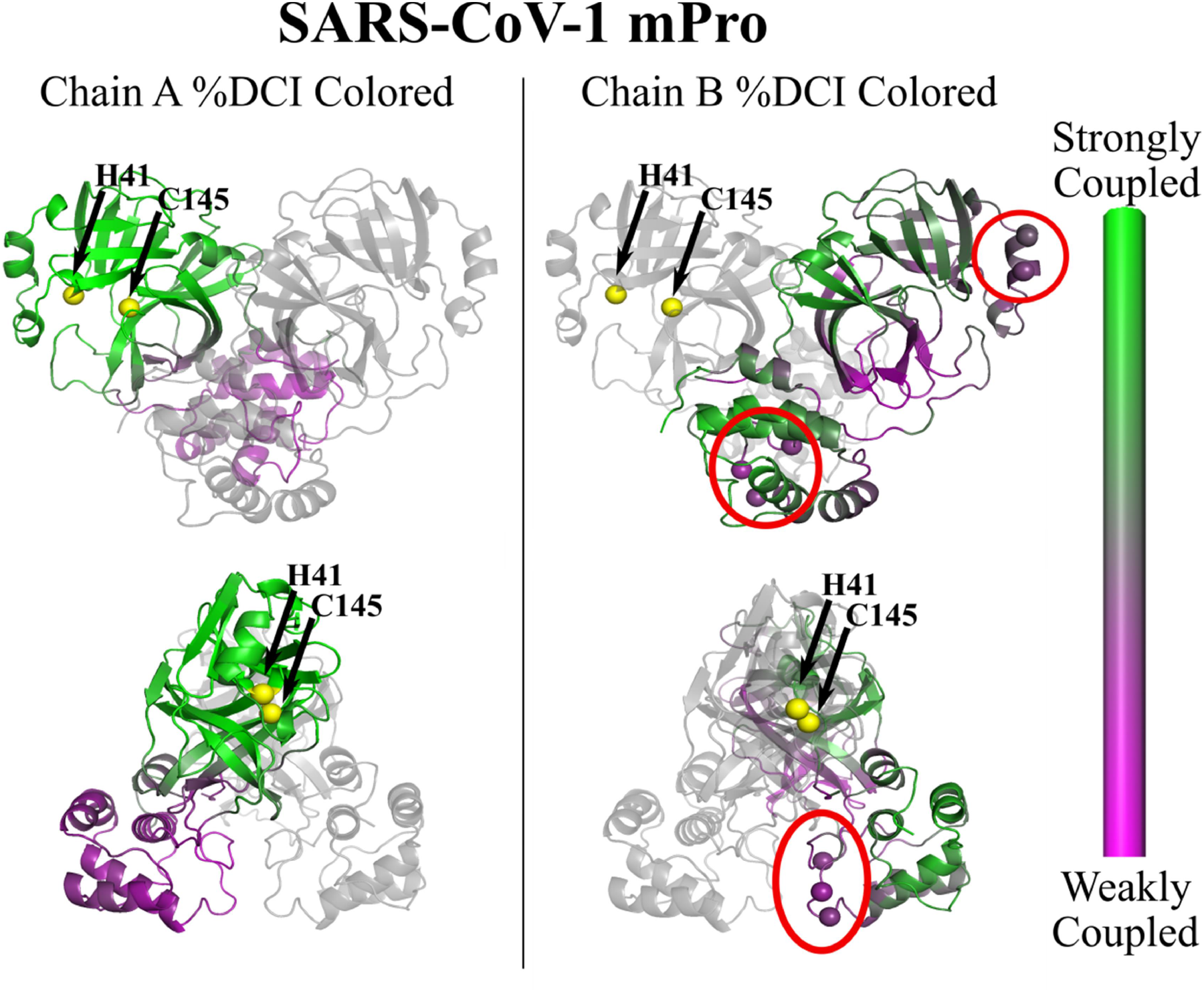
Dynamic coupling analysis (measured by %DCI) between catalytic site residues of H41 and C145 (yellow spheres) of chain A and the other residues of chain A (left) and chain B (right) mapped onto the SARS-CoV-1 mPro (PDB ID 2GZ9(32), inhibitors removed). Green regions of each structure indicate stronger dynamic coupling while purple represents weak dynamic coupling. Within the chain (left), the %DCI distribution is expected based off proximity; residues close to the catalytic sites are strongly coupled, with those further away exhibiting weak coupling and exhibits much the same behavior as the SARS-CoV-2 mPro. In cross-chain coupling to chain B (right), this behavior is maintained, opposite to that of SARS-CoV-2 (Figure 1). Chain B sites which exhibited particularly strong coupling to the chain A catalytic residues in the CoV-2 mPro are circled in red and shown as spheres; here we see the strong coupling of the catalytic sites to these residues is absent in CoV-1.

Given the noticeable differences in coupling between the two systems, we next wanted to compare the changes in dynamic coupling to the active site residues of the mutation sites themselves. Figure 3 shows this direct comparison. In Figure 3 (A) we have taken the %DCI values from the chain A active site perturbations shown in Figures 1 and 2 and subtracted the CoV-2 %DCI profile from the CoV-1 profile. Here, all sites with a difference in %DCI within one standard deviation of the mean have been set to zero .This analysis immediately shows that, indeed, sites which exhibit large changes in dynamic coupling are often not at the sites of mutation themselves. The %DCI values used for this comparison are plotted in Figure 3 (B) with dashed lines indicating the sites of mutation. Interestingly, we see notable changes in coupling in areas surrounding the active sites of *both* chains, even though only the active site residues from chain A were perturbed. Particularly, we see enhanced coupling to the loop residues 185-201 (marked in black ovals), an area previously reported to assist in stabilizing the active sites (33).

Additionally, recent work has been performed to identify positions within the SARS-CoV-2 mPro resistant to mutations in human populations. From a dataset containing 19,154 mutations it was shown that 282 out of 306 residue positions of SARS-CoV-2 have exhibited at least one mutation (34); excluding the active sites 41 and 145, the 24 remaining positions which had not experienced any mutation events were deemed “coldspots” and subjected to further study (35). Interestingly, a comparative analysis of the change in flexibility of these coldspots (of both chains A and B) between CoV-2 and CoV-1 shows that, while evolutionarily conserved, these sites experienced a significantly greater change in dynamic flexibility (as measured by %DFI) when compared to all residues in the structure (see supplemental figure S1). Additionally, performing dynamic coupling analysis of these coldspots residues uncovered a unique relationship for six specific positions. Through crystallographic studies, positions Leu141, Phe185 and Gln192 of both chains A and B were shown to be structurally important, involved in the formation of substrate-binding sites (36–38). When we analyzed these positions’ dynamic coupling to catalytic site residues His41 and Cys145, we found that while the intrachain coupling remained relatively unchanged (within one standard deviation of the mean), the cross-chain dynamic coupling was significantly enchanced in SARS-CoV-2 with all three sites showing an increase in %DCI greater than one standard deviation from the mean. Further, positions Phe185 and Gln192, which play critical roles in stabilizing the active sites (36–38), exhibited %DCI increases greater than two standard deviations from the mean.. Since many of the large changes in coupling are located at sites distal to the mutation sites, we wanted to next understand the changes between the two systems via site-specific flexibility as these changes in coupling could be due to dynamical differences which arise from changes in amino acid flexibility.

**Figure 3.**
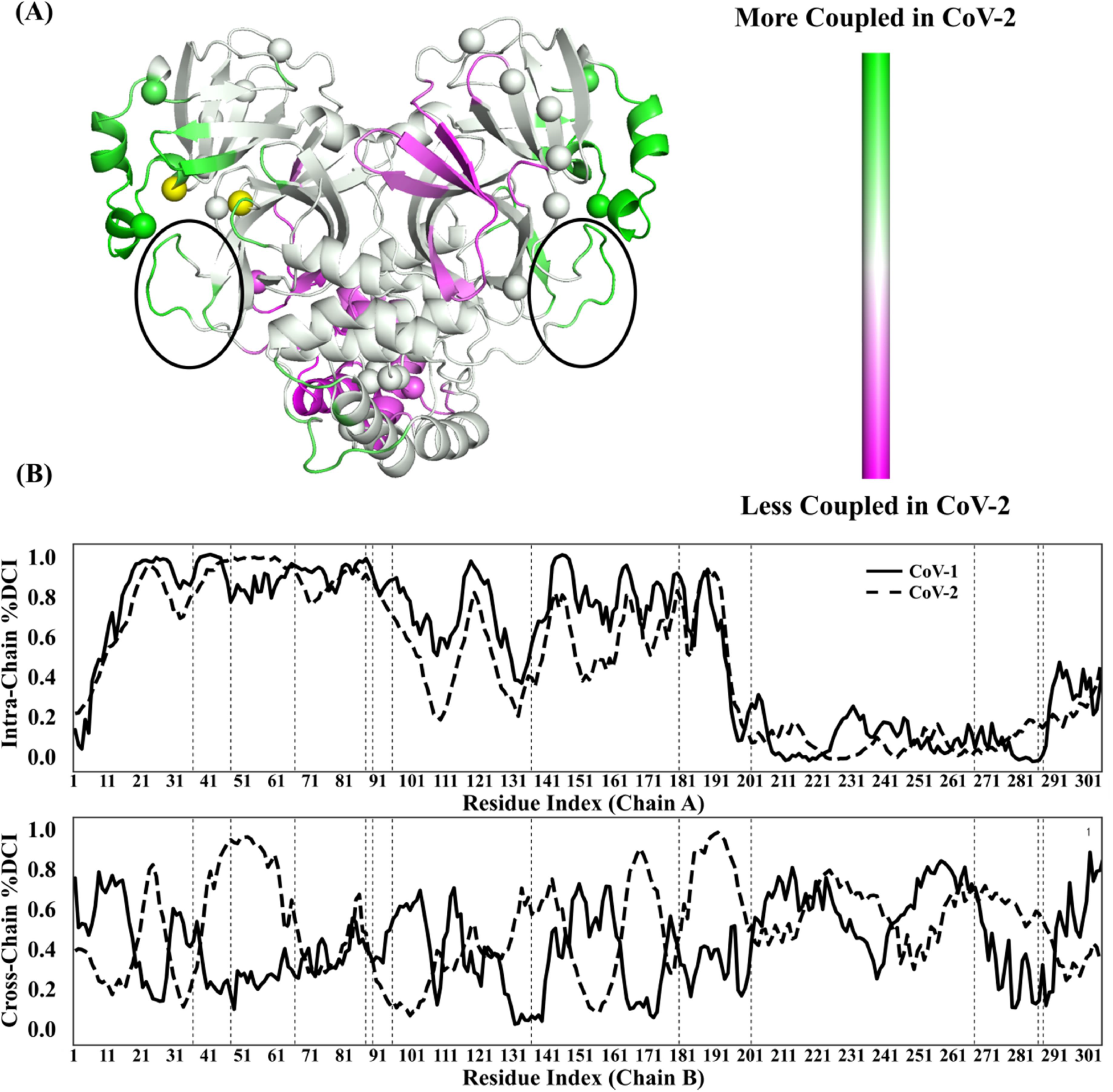
**(A)** %DCI difference between the two systems (CoV-2 %DCI – CoV-1 %DCI) mapped onto the CoV-2 (PDB ID 5R7Y(31)) main protease using MD covariance matrix data to generate the profiles. The active sites perturbed to generate these values are marked in yellow spheres, as in Figures 1 and 2. Here, all sites with a difference in %DCI within one standard deviation of the mean have been set to zero, thus, purple indicates a significant loss in dynamic coupling whereas green indicates a significant gain in dynamic coupling in COV-2. Sites of mutation (positions 35,46,65,86,88,94,134,180,202,267,285,286) are also shown as spheres. **(B)** %DCI values used for the subtraction in **(A)** where dashed lines mark mutation sites. In most cases, the mutation sites themselves are not those exhibiting the greatest change in coupling to the active sites.

### Dynamical differences between the CoV-1 and CoV-2 proteases occur at sites distal to the mutational sites themselves

In an effort to understand the mechanics underlying the behavior of the SARS-CoV-2 mPro, a comparative analysis of the dynamical differences between SARS-CoV-1 and SARS-CoV-2 was performed to capture any major differences in flexibility. Particularly, we wanted to investigate the dynamic flexibility differences as related to inhibitory binding events. To this end we studied the mPro of each virus using the Dynamic Flexibility Index (DFI). Similar to DCI, the dynamic DFI combines PRS and LRT(26,23) to evaluate each position’s displacement response to random force perturbations at other locations in the protein. For each position *i*, the DFI value quantifies its total displacement response relative to the total displacement response for all positions in the protein. To identify the most flexible positions, ranked *%*DFI values are used: For each position *i*, its percentile rank in the observed DFI range is calculated from the ratio of the number of positions with DFI values ≤ DFI_i_ to the total number of positions. *%*DFI values range from 0 to 1. The DFI parameter can be considered a measure of a given position’s ability to explore its local conformational space; that is, DFI is a measure of an amino acid’s flexibility and, in the case of homodimeric systems, is largely symmetric across monomeric units. This parameter has been used to identify important structural elements of a protein, such as hinges, and to show the flexibility of functionally critical areas, such as binding domains and catalytic regions. Surprisingly, when the flexibility profiles of the two structures were compared, we found that, on average, the larger changes in dynamic flexibility were occurring at positions *other than* the mutation sites themselves (only ∼20% of mutated sites experienced a %DFI difference outside of one standard deviation, whereas 31% of all other residues fell into this category). (Fig. 4). Also, of interesting note, the loop regions 185-201, previously mentioned above as mechanistically important for stabilizing the active sites, does not exhibit a large change in flexibility in the CoV-2 system, also in line with previous work (33).

**Figure 4.**
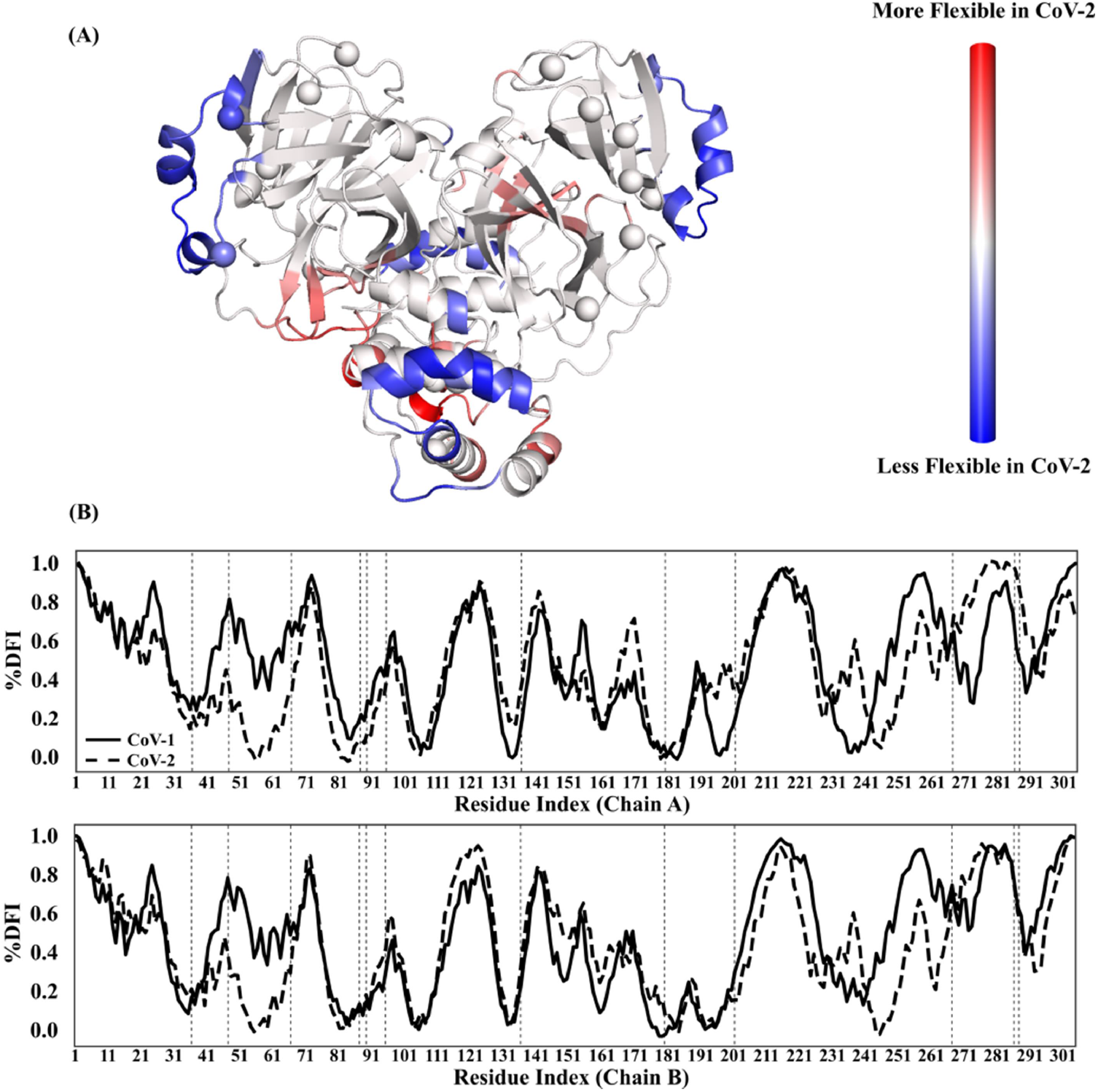
**(A)** Chain A flexibility comparison between SARS-CoV-1 CoV-2 measured by the difference in %DFI (CoV-2 %DFI – CoV-1 %DFI) mapped onto the CoV-2 mPro (PDB ID 5R7Y(31)) using MD covariance matrices where red indicates a gain in flexibility while blue indicates a loss in flexibility. Here, all sites with a difference in %DFI within one standard deviation of the mean have been set to zero to identify the significant flexibility changes in COV-2.. The sites of mutation between the two structures (positions 35,46,65,86,88,94,134,180,202,267,285,286) are marked in spheres. **(B)** The %DFI values used to generate the color-coding in **(A)** where mutation sites are marked by dashed lines. Of note, many of the sites which exhibit large differences in %DFI occur at positions other than the mutation sites themselves.

Of specific interest is the identification of sites exhibiting extremely flow flexibility (marked by %DFI values <= 0.2). While positions with low flexibility are often evolutionarily conserved (39–42), our %DFI analysis shows the formation of large regions of low flexibility spanning residues 51-63 of both chains A and B in the SARS-CoV-2, absent in the CoV-1 system. These low flexibility sites called “hinges” have previously been shown to be mechanistically critical in determining or regulating protein function (43,44,28). Hinges play a pivotal role in the transfer of force through external perturbations throughout the chain in a cascading fashion and often control and mediate protein motion, similar to joints in a skeleton. Further, the formation of hinges, or, “hinge-shift mechanisms”, have been linked to gain-in-function in multiple different enzyme families via protein evolution (45,24,46,47). These results indicate that the biophysical differences between the two systems may be explained, mechanically, through allosteric modulation. Here, the mutations cause changes in flexibility of amino acid positions located elsewhere in the structure and, subsequently, result in differing dynamics between the two structures.

### Inhibitor Binding Allosterically Induces Changes in Flexibility to Distal Sites Differently between CoV-1 and CoV-2

As comparative flexibility analysis above suggests that allosteric regulation may be a key component in capturing the changes in dynamics between the CoV-1 and CoV-2 mPro systems, we next wanted to determine how the allosteric response to ligand binding events (particularly inhibitor binding) differed between the two. In fact, recent work using Gaussian accelerated molecular dynamics indicates that there are potential cryptic binding pockets located far from the active site which may act to inhibit the active sites allosterically when bound to inhibitory drugs (48). To further analyze the allosteric impact of inhibitor binding, we employed the Elastic Network Model (ENM) formalism for DFI (see: Methods) which allows for rapid modeling of ligand binding events by adding additional alpha carbon atoms at the inhibitor binding sites, thus extending each protein’s residue-residue anisotropic network to include modeled inhibitors. For CoV-1, the main protease complexed with an alpha, beta-unsaturated ethyl ester inhibitor SG82 (PDB ID 3TIU) was chosen to be inhibitor bound form. Additionally, we used main protease in complex with carmofur (PDB ID 7BUY) as the inhibitor bound form for CoV-2. Both inhibitors are bound at the catalytic site, and four carbon atoms among each inhibitor were picked as alpha carbon atoms which contribute as nodes in ENM calculation.

Figure 5 shows the %DFI values mapped onto the three dimensional structures of each mPro. While, overall, the dynamic flexibility profiles between structures were similar in both inhibitor-unbound (top) and inhibitor-bound (bottom), a specific difference in behavior was observed for a set of residues (N277, R279 and L286, black circle) located at the dimeric interface of both chains A and B. Notably, when bound to an inhibitor, these residues of the CoV-1 mPro became *more* flexible, whereas the same residues in the inhibitor-bound CoV-2 mPro became *less* flexible. Interestingly, normal mode analysis utilizing ENM performed by suggests that the first mode of the dimeric protease involves the opening / closing of this region of the protein (49) which may be altered significantly upon a ligand binding event, particularly if bound in the dimeric form. These results indicate that not only does an inhibitor binding event at the catalytic site residues induce flexibility changes at regions distal to the binding site, but does so in a manner opposite between the CoV-1 and CoV-2 mPro, with the residues N277, R279 and L286 becoming more / less flexible, respectively. As such a binding event allows for allosteric regulation of the protein, it follows that there is likely additional allosteric regulation between the important catalytic site residues H41 and C145 with the rest of the protein.

**Figure 5.**
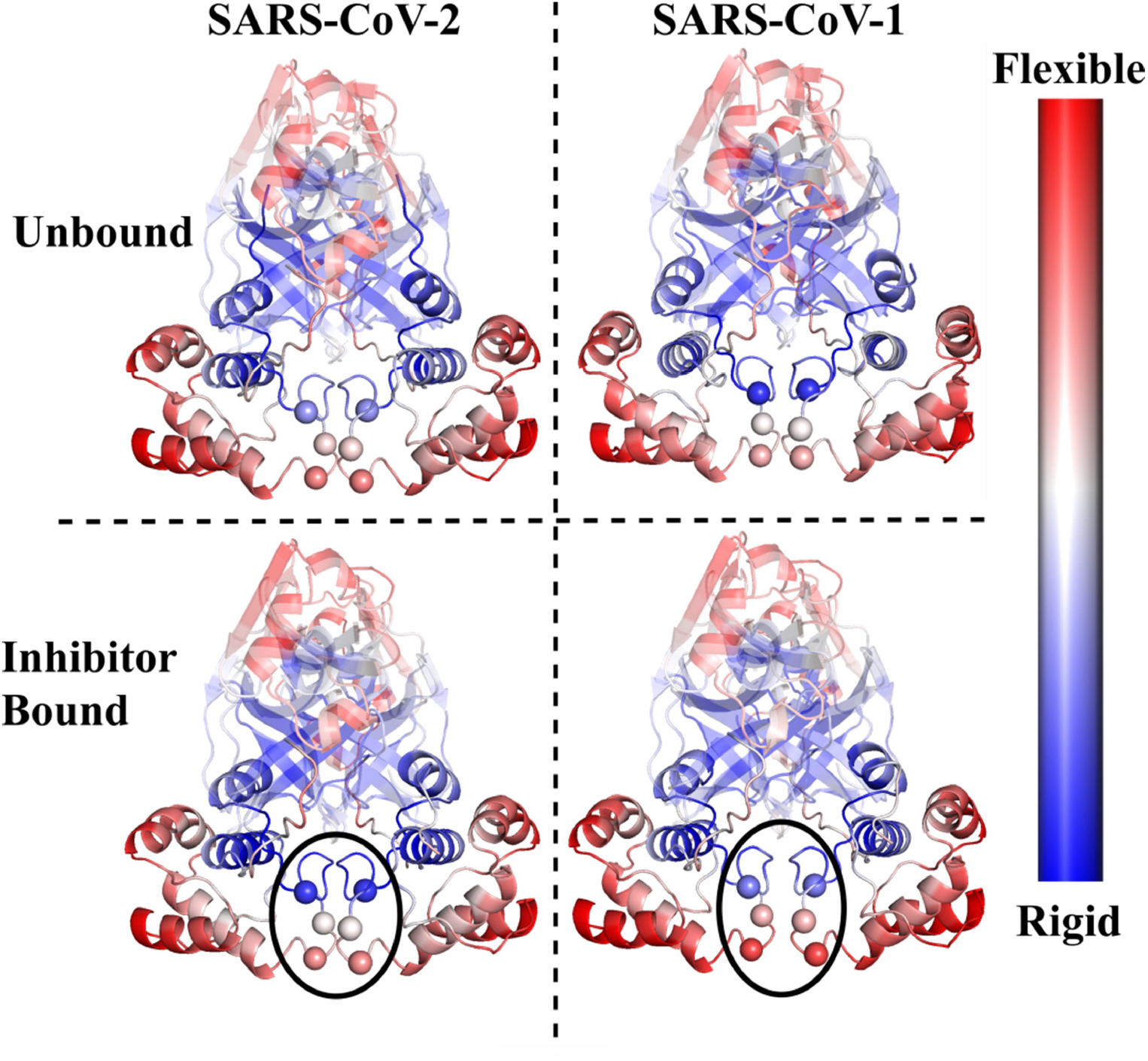
Comparison of dynamic flexibility (measured by %DFI) mapped onto the inhibitor bound (top) and inhibitor unbound (bottom) forms of the SARS-CoV-2 (left) PDB ID 6M03(37)(unbound) and PDB ID 7BUY(17) (bound) and SARS-CoV-1 (right) PDB ID 1UK3(50)(unbound) and PDB ID 3TIU(51) (bound). Generally, the dynamic flexibility profiles between structures were similar in both inhibitor-unbound and inhibitor-bound (bottom). A specific difference in behavior was observed for a set of residues (N277, R279 and L286, black circle) located at the dimeric interface of both chains A and B. Notably, when bound to an inhibitor, these residues of the CoV-1 mPro became *more* flexible, whereas the same residues in the inhibitor-bound CoV-2 mPro became *less* flexible. These figures were rendered using PyMOL(52).

## CONCLUSION

Here we have conducted a comparative analysis between the SARS-CoV-1 and SARS-CoV-2 mPro systems to shed mechanistic insight on the biophysical changes associated with the mutations between these two enzymes. Our work shows that the CoV-2 mPro system exhibits enhanced cross-chain communication between catalytic site residues and the rest of the structure. Further, both dynamic coupling and dynamic flexibility analyses indicates that, largely, the dynamic changes as evaluated by DCI and DFI occur at sites *other than* the mutation sites themselves, indicating that the functional differences between these two proteins are a result of dynamic allostery induced elsewhere in the structure upon these mutations. Finally, we show that specific regions of the CoV-2 mPro exhibit markedly different flexibility behavior when bound to an inhibitior as compared to the CoV-1 mPro. Future work will focus on the analysis of these signature regions with the greatest change in coupling and flexibility in an effort to identify putative allosteric binding sites for potential inhibitory drug discovery.

## AUTHOR CONTRIBUTIONS

P. Campitelli and J. Lu each generated molecular dynamics data for the CoV-1 and CoV-2 systems. P. Campitelli conducted all analysis presented here with the exception of the ENM modeled, ligand-bound work which was performed by J. Lu. P. Campitelli wrote the manuscript with additional writing contributions and editing from both J. Lu and S. B. Ozkan. S. B. Ozkan was responsible for the oversight and narrative of the analysis found in this manuscript.

## ACKNOWLEDGEMENTS

This work is supported by the National Science Foundation Division of Molecular and Cellular Biosciences (award 1715591) and the Gorden and Betty Moore Foundation Award #8415

## REFERENCES

1. U.S. Food and Drug Administration. PFIZER-BIONTECH COVID-19 VACCINE. https://www.fda.gov/emergency-preparedness-and-response/coronavirus-disease-2019-covid-19/pfizer-bion-tech-covid-19-vaccine. Accessed March 08, 2021.

2. U.S. Food and Drug Administration. MODERNA COVID-19 VACCINE. https://www.fda.gov/emergency-preparedness-and-response/coronavirus-disease-2019-covid-19/moderna-covid-19-vaccine. Accessed March 08, 2021.

3. Polack, F. P., S. J. Thomas, N. Kitchin, J. Absalon, and A. Gurtman et al. 2020. SAFETY AND EFFICACY OF THE BNT162B2 MRNA COVID-19 VACCINE. The New England journal of medicine, 383:2603–2615.

4. Jackson, L. A., E. J. Anderson, N. G. Rouphael, P. C. Roberts, and M. Makhene et al. 2020. AN MRNA VACCINE AGAINST SARS-COV-2 - PRELIMINARY REPORT. The New England journal of medicine, 383:1920–1931.

5. Kahn, B., L. Brown, W. Foege, and H. Gayle. 2020. FRAMEWORK FOR EQUITABLE ALLOCATION OF COVID-19 VACCINE, Washington (DC).

6. Khubchandani, J., S. Sharma, J. H. Price, M. J. Wiblishauser, and M. Sharma et al. 2021. COVID-19 VACCINATION HESITANCY IN THE UNITED STATES: A RAPID NATIONAL ASSESSMENT. Journal of community health, 46:270–277.

7. McAteer, J., I. Yildirim, and A. Chahroudi. 2020. THE VACCINES ACT: DECIPHERING VACCINE HESITANCY IN THE TIME OF COVID-19. Clinical infectious diseases : an official publication of the Infectious Diseases Society of America, 71:703–705.

8. Ball, P. 2020. ANTI-VACCINE MOVEMENT COULD UNDERMINE EFFORTS TO END CORONAVIRUS PANDEMIC, RESEARCHERS WARN. Nature, 581:251.

9. Hunter, P. R., J. Brainard, and A. Grant. 2021. THE IMPACT OF THE NOVEMBER 2020 ENGLISH NATIONAL LOCKDOWN ON COVID-19 CASE COUNTS. medRxiv.

10. Volz, E., S. Mishra, M. Chand, J. C. Barrett, and R. Johnson et al. 2021. TRANSMISSION OF SARS-COV-2 LINEAGE B.1.1.7 IN ENGLAND: INSIGHTS FROM LINKING EPIDEMIOLOGICAL AND GENETIC DATA. medRxiv.

11. Tegally, H., E. Wilkinson, R. J. Lessells, J. Giandhari, and S. Pillay et al. 2021. SIXTEEN NOVEL LINEAGES OF SARS-COV-2 IN SOUTH AFRICA. Nature medicine, 27:440–446.

12. Washington, N. L., K. Gangavarapu, M. Zeller, A. Bolze, and E. T. Cirulli et al. 2021. EMERGENCE AND RAPID TRANSMISSION OF SARS-COV-2 B.1.1.7 IN THE UNITED STATES. Cell, 184:2587-2594.e7.

13. Ren, Z., L. Yan, N. Zhang, Y. Guo, and C. Yang et al. 2013. THE NEWLY EMERGED SARS-LIKE CORONAVIRUS HCOV-EMC ALSO HAS AN “ACHILLES’ HEEL”: CURRENT EFFECTIVE INHIBITOR TARGETING A 3C-LIKE PROTEASE. Protein & cell, 4:248–250.

14. Ramajayam, R., K.-P. Tan, and P.-H. Liang. 2011. RECENT DEVELOPMENT OF 3C AND 3CL PROTEASE INHIBITORS FOR ANTI-CORONAVIRUS AND ANTI-PICORNAVIRUS DRUG DISCOVERY. Biochemical Society transactions, 39:1371–1375.

15. Dai, W., B. Zhang, X.-M. Jiang, H. Su, and J. Li et al. 2020. STRUCTURE-BASED DESIGN OF ANTIVIRAL DRUG CANDIDATES TARGETING THE SARS-COV-2 MAIN PROTEASE. Science, 368:1331–1335.

16. Das, S., S. Sarmah, S. Lyndem, and A. Singha Roy. 2020. AN INVESTIGATION INTO THE IDENTIFICATION OF POTENTIAL INHIBITORS OF SARS-COV-2 MAIN PROTEASE USING MOLECULAR DOCKING STUDY. Journal of biomolecular structure & dynamics, 39:3347–3357.

17. Jin, Z., Y. Zhao, Y. Sun, B. Zhang, and H. Wang et al. 2020. STRUCTURAL BASIS FOR THE INHIBITION OF SARS-COV-2 MAIN PROTEASE BY ANTINEOPLASTIC DRUG CARMOFUR. Nature structural & molecular biology, 27:529–532.

18. Ullrich, S., and C. Nitsche. 2020. THE SARS-COV-2 MAIN PROTEASE AS DRUG TARGET. Bioorganic & medicinal chemistry letters, 30:127377.

19. Tekpinar, M., and A. Yildirim. 2021. IMPACT OF DIMERIZATION AND N3 BINDING ON MOLECULAR DYNAMICS OF SARS-COV AND SARS-COV-2 MAIN PROTEASES. Journal of biomolecular structure & dynamics:1–12.

20. Sheik Amamuddy, O., G. M. Verkhivker, and Ö. Tastan Bishop. 2020. IMPACT OF EARLY PANDEMIC STAGE MUTATIONS ON MOLECULAR DYNAMICS OF SARS-COV-2 MPRO. Journal of chemical information and modeling, 60:5080–5102.

21. Suárez, D., and N. Díaz. 2020. SARS-COV-2 MAIN PROTEASE: A MOLECULAR DYNAMICS STUDY. Journal of chemical information and modeling, 60:5815–5831.

22. McLeish, T. C., and I. Dubanevics. 2021. COMPUTATIONAL ANALYSIS OF DYNAMIC ALLOSTERY AND CONTROL IN THE SARS-COV-2 MAIN PROTEASE. Journal of the Royal Society, Interface, 18.

23. Nevin Gerek, Z., S. Kumar, and S. Banu Ozkan. 2013. STRUCTURAL DYNAMICS FLEXIBILITY INFORMS FUNCTION AND EVOLUTION AT A PROTEOME SCALE. Evolutionary Applications, 6:423–433.

24. Modi, T., J. Huihui, K. Ghosh, and S. B. Ozkan. 2018. ANCIENT THIOREDOXINS EVOLVED TO MODERN-DAY STABILITY-FUNCTION REQUIREMENT BY ALTERING NATIVE STATE ENSEMBLE. Philosophical transactions of the Royal Society of London. Series B, Biological sciences, 373.

25. Modi, T., and S. B. Ozkan. 2018. MUTATIONS UTILIZE DYNAMIC ALLOSTERY TO CONFER RESISTANCE IN TEM-1 Β-LACTAMASE. International Journal of Molecular Sciences, 19.

26. Gerek, Z. N., and S. B. Ozkan. 2011. CHANGE IN ALLOSTERIC NETWORK AFFECTS BINDING AFFINITIES OF PDZ DOMAINS: ANALYSIS THROUGH PERTURBATION RESPONSE SCANNING. PLoS Computational Biology, 7.

27. Campitelli, P., L. Swint-Kruse, and S. B. Ozkan. 2020. SUBSTITUTIONS AT NON-CONSERVED RHEOSTAT POSITIONS MODULATE FUNCTION BY RE-WIRING LONG-RANGE, DYNAMIC INTERACTIONS. Molecular biology and evolution, 38:201–214.

28. Campitelli, P., J. Guo, H.-X. Zhou, and S. B. Ozkan. 2018. HINGE-SHIFT MECHANISM MODULATES ALLOSTERIC REGULATIONS IN HUMAN PIN1. The journal of physical chemistry. B, 122:5623–5629.

29. Kumar, A., B. M. Butler, S. Kumar, and S. B. Ozkan. 2015. INTEGRATION OF STRUCTURAL DYNAMICS AND MOLECULAR EVOLUTION VIA PROTEIN INTERACTION NETWORKS: A NEW ERA IN GENOMIC MEDICINE. Current opinion in structural biology, 35:135–142.

30. Case, D. A., T. E. Cheatham, T. Darden, H. Gohlke, and R. Luo et al. 2005. THE AMBER BIOMOLECULAR SIMULATION PROGRAMS. Journal of computational chemistry, 26:1668–1688.

31. Douangamath, A., D. Fearon, P. Gehrtz, T. Krojer, and P. Lukacik et al. 2020. CRYSTALLOGRAPHIC AND ELECTROPHILIC FRAGMENT SCREENING OF THE SARS-COV-2 MAIN PROTEASE. Nature communications, 11:5047.

32. Lu, I.-L., N. Mahindroo, P.-H. Liang, Y.-H. Peng, and C.-J. Kuo et al. 2006. STRUCTURE-BASED DRUG DESIGN AND STRUCTURAL BIOLOGY STUDY OF NOVEL NONPEPTIDE INHIBITORS OF SEVERE ACUTE RESPIRATORY SYNDROME CORONAVIRUS MAIN PROTEASE. Journal of medicinal chemistry, 49:5154–5161.

33. Weng, Y. L., S. R. Naik, N. Dingelstad, M. R. Lugo, and S. Kalyaanamoorthy et al. 2021. MOLECULAR DYNAMICS AND IN SILICO MUTAGENESIS ON THE REVERSIBLE INHIBITOR-BOUND SARS-COV-2 MAIN PROTEASE COMPLEXES REVEAL THE ROLE OF LATERAL POCKET IN ENHANCING THE LIGAND AFFINITY. Scientific Reports, 11:7429.

34. Badua, C. L. D. C., K. A. T. Baldo, and P. M. B. Medina. 2021. GENOMIC AND PROTEOMIC MUTATION LANDSCAPES OF SARS-COV-2. J Med Virol, 93:1702–1721.

35. Krishnamoorthy, N., and K. Fakhro. 2021. IDENTIFICATION OF MUTATION RESISTANCE COLDSPOTS FOR TARGETING THE SARS-COV2 MAIN PROTEASE. IUBMB life, 73:670–675.

36. Douangamath, A., D. Fearon, P. Gehrtz, T. Krojer, and P. Lukacik et al. 2020. CRYSTALLOGRAPHIC AND ELECTROPHILIC FRAGMENT SCREENING OF THE SARS-COV-2 MAIN PROTEASE. Nat Commun, 11:5047.

37. Jin, Z., X. Du, Y. Xu, Y. Deng, and M. Liu et al. 2020. STRUCTURE OF MPRO FROM SARS-COV-2 AND DISCOVERY OF ITS INHIBITORS. Nature, 582:289–293.

38. Kneller, D. W., G. Phillips, K. L. Weiss, S. Pant, and Q. Zhang et al. 2020. UNUSUAL ZWITTERIONIC CATALYTIC SITE OF SARS-COV-2 MAIN PROTEASE REVEALED BY NEUTRON CRYSTALLOGRAPHY. The Journal of biological chemistry, 295:17365–17373.

39. Liu, Y., and I. Bahar. 2012. SEQUENCE EVOLUTION CORRELATES WITH STRUCTURAL DYNAMICS. Molecular biology and evolution, 29:2253–2263.

40. Maguid, S., S. Fernandez-Alberti, and J. Echave. 2008. EVOLUTIONARY CONSERVATION OF PROTEIN VIBRATIONAL DYNAMICS. Gene, 422:7–13.

41. Maguid, S., S. Fernández-Alberti, G. Parisi, and J. Echave. 2006. EVOLUTIONARY CONSERVATION OF PROTEIN BACKBONE FLEXIBILITY. Journal of molecular evolution, 63:448–457.

42. Miller, M., Y. Bromberg, and L. Swint-Kruse. 2017. COMPUTATIONAL PREDICTORS FAIL TO IDENTIFY AMINO ACID SUBSTITUTION EFFECTS AT RHEOSTAT POSITIONS. Scientific Reports, 7:41329.

43. Kumar, A., T. J. Glembo, and S. B. Ozkan. 2015. THE ROLE OF CONFORMATIONAL DYNAMICS AND ALLOSTERY IN THE DISEASE DEVELOPMENT OF HUMAN FERRITIN. Biophysical Journal, 109:1273–1281.

44. Li, Z., A. Bolia, J. D. Maxwell, A. A. Bobkov, and G. Ghirlanda et al. 2015. A RIGID HINGE REGION IS NECESSARY FOR HIGH-AFFINITY BINDING OF DIMANNOSE TO CYANOVIRIN AND ASSOCIATED CONSTRUCTS. Biochemistry, 54:6951–6960.

45. Kim, H., T. Zou, C. Modi, K. Dörner, and T. J. Grunkemeyer et al. 2015. A HINGE MIGRATION MECHANISM UNLOCKS THE EVOLUTION OF GREEN-TO-RED PHOTOCONVERSION IN GFP-LIKE PROTEINS. Structure (London, England : 1993), 23:34–43.

46. Zou, T., V. A. Risso, J. A. Gavira, J. M. Sanchez-Ruiz, and S. B. Ozkan. 2015. EVOLUTION OF CONFORMATIONAL DYNAMICS DETERMINES THE CONVERSION OF A PROMISCUOUS GENERALIST INTO A SPECIALIST ENZYME. Molecular biology and evolution, 32:132–143.

47. Zheng, W., B. R. Brooks, S. Doniach, and d. Thirumalai. 2005. NETWORK OF DYNAMICALLY IMPORTANT RESIDUES IN THE OPEN/CLOSED TRANSITION IN POLYMERASES IS STRONGLY CONSERVED. Structure (London, England : 1993), 13:565–577.

48. Sztain, T., R. Amaro, and J. A. McCammon. 2021. ELUCIDATION OF CRYPTIC AND ALLOSTERIC POCKETS WITHIN THE SARS-COV-2 MAIN PROTEASE. Journal of chemical information and modeling.

49. Chen, Y. W., C.-P. B. Yiu, and K.-Y. Wong. 2020. PREDICTION OF THE SARS-COV-2 (2019-NCOV) 3C-LIKE PROTEASE (3CL PRO) STRUCTURE: VIRTUAL SCREENING REVEALS VELPATASVIR, LEDIPASVIR, AND OTHER DRUG REPURPOSING CANDIDATES. F1000Research, 9:129.

50. Yang, H., M. Yang, Y. Ding, Y. Liu, and Z. Lou et al. 2003. THE CRYSTAL STRUCTURES OF SEVERE ACUTE RESPIRATORY SYNDROME VIRUS MAIN PROTEASE AND ITS COMPLEX WITH AN INHIBITOR. Proc Natl Acad Sci USA, 100:13190–13195.

51. Zhu, L., and R. Hilgenfeld. 2012. CRYSTAL STRUCTURE OF SARS CORONAVIRUS MAIN PROTEASE COMPLEXED WITH AN ALPHA,BETA-UNSATURATED ETHYL ESTER INHIBITOR SG82.

52. THE PYMOL MOLECULAR GRAPHICS SYSTEM (Schrodinger LLC).

53. MacKerell, A. D., D. Bashford, M. Bellott, R. L. Dunbrack, and J. D. Evanseck et al. 1998. ALL-ATOM EMPIRICAL POTENTIAL FOR MOLECULAR MODELING AND DYNAMICS STUDIES OF PROTEINS. The journal of physical chemistry. B, 102:3586–3616.

54. Darden, T., D. York, and L. Pedersen. 1993. PARTICLE MESH EWALD: AN N LOG(N) METHOD FOR EWALD SUMS IN LARGE SYSTEMS. The EMBO journal, 98:10089–10092.

