## Supplemental figure 1 for "Dynamic Allostery Highlights the Evolutionary Differences between the CoV-1 and CoV-2 Main Proteases"

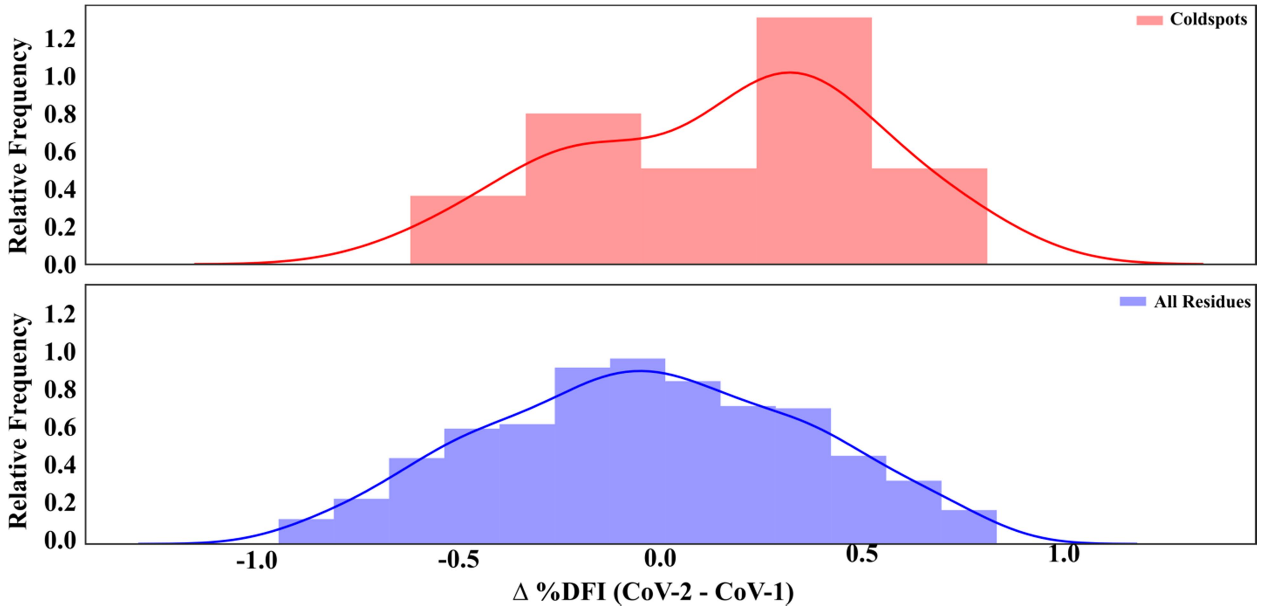


**Figure S1.** Comparison of change in flexibility (as measured by %DFI) between the SARS-CoV-2 and CoV-1 mPros of "coldspots" (i.e. residues which exhibited no mutations between the two systems within the human population) versus all residues within the structure. Interestingly, while evolutionarily conserved, these positions exhibited a greater-than-average change in flexibilty, with many of them becoming more flexibile in CoV-2.
